# A genome-wide haploid genetic screen for essential factors in vaccinia virus infection identifies TMED10 as regulator of macropinocytosis

**DOI:** 10.1101/493205

**Authors:** Rutger D. Luteijn, Ferdy van Diemen, Vincent A. Blomen, Ingrid G.J. Boer, Saran Manikam Sadasivam, Toin H. van Kuppevelt, Ingo Drexler, Thijn R. Brummelkamp, Robert Jan Lebbink, Emmanuel J. Wiertz

**Author notes:** These authors contributed equally to this paper (Robert Jan Lebbink, Emmanuel J. Wiertz).

## Abstract

Vaccinia virus is a promising viral vaccine and gene delivery candidate, and has historically been used as a model to study poxvirus-host cell interactions. We employed a genome-wide insertional mutagenesis approach in human haploid cells to identify host factors crucial for vaccinia virus infection. A library of mutagenized HAP1 cells was exposed to Modified Vaccinia Virus Ankara (MVA). Deep-sequencing analysis of virus-resistant cells identified host factors involved in heparan sulfate synthesis, Golgi organization, and vesicular protein trafficking. We validated EXT1, TM9SF2 and TMED10 (TMP21/p23/p24δ) as important host factors for vaccinia virus infection. The critical role of EXT1 in heparan sulfate synthesis and vaccinia virus infection was confirmed. TM9SF2 was validated as a player mediating heparan sulfate expression, explaining its contribution to vaccinia virus infection. In addition, TMED10 was found to be crucial for virus-induced plasma membrane blebbing and phosphatidylserine-induced macropinocytosis, suggesting that TMED10 regulates actin cytoskeleton remodelling necessary for virus infection.

**Importance:** Poxviruses are large DNA viruses that can infect a wide range of host species. A number of these viruses are clinically important to humans, including variola virus (smallpox) and vaccinia virus. Since the eradication of smallpox, zoonotic infections with monkeypox virus and cowpox virus are emerging. Additionally, poxviruses can be engineered to specifically target cancer cells, and are used as vaccine vector against tuberculosis, influenza, and coronaviruses.

Poxviruses rely on host factors for most stages of their life cycle, including attachment to the cell and entry. These host factors are crucial for virus infectivity and host cell tropism. We used a genome-wide knock-out library of host cells to identify host factors necessary for vaccinia virus infection. We confirm a dominant role for heparin sulfate in mediating virus attachment. Additionally, we show that TMED10, previously not implicated in virus infections, modulates the host cell membrane to facilitate virus uptake.

## Introduction

The poxvirus family represents a group of large enveloped DNA viruses that infect a wide variety of hosts. Poxvirus species capable of infecting humans include variola virus, which causes smallpox and is one of the most destructive pathogens in human history. Since its eradication through successful vaccination using vaccinia virus, poxvirus outbreaks in humans are nowadays mainly caused by zoonotic infections of cowpox virus, monkeypox virus, and recently discovered poxvirus species (1-3). The number of zoonotic infections is predicted to rise, due to waning population immunity (4). In part, the decreased immunity is caused by concerns about vaccinia virus safety, as a minority of vaccinated individuals show adverse side effects (5). These safety concerns have led to the development of safer, attenuated vaccinia virus strains, including the Modified Vaccinia Ankara virus (MVA). By passaging vaccinia virus over 500 times on chicken embryo fibroblasts, the resulting attenuated MVA lost 10% of the parental vaccinia genome, and displays an abortive replication cycle in most cell lines (6). MVA is potent vaccine against poxviruses and serves as a vaccine vector against a variety of other diseases (7-13).

Due to their low pathogenicity and wide range of applications, vaccinia virus strains are used as model viruses to study the unique life cycle of poxviruses (14). The vaccinia virus life cycle starts with binding to heparan sulfate (HepS) and other glycosaminoglycans expressed on the host cell, although laminin and unidentified cellular proteins may also play a role (15). Upon binding, the viral envelope fuses with the host cell at either the plasma membrane or the endosomal membrane after macropinocytotic uptake by the host (16). Release of the virus core in the cytoplasm initiates transcription of more than 100 viral genes (17). Early gene transcripts mediate virus core uncoating and DNA replication (18). Replicated viral genomes serve as template for transcription of intermediate and late genes, many of which are involved in the assembly of virus progeny (18).

In contrast to most DNA viruses, poxvirus replication is located outside the cellular nucleus in specialized compartments known as viral factories (14). Due to their independence from the host nucleus, poxviruses are required to encode most of the genes involved in DNA replication and transcription. Consequently, poxviruses are considered to be less dependent on host factors compared to other DNA viruses. Nevertheless, interaction with the host cell are required during most stages of virus infection. Here, we employed a genome-wide haploid genetic screen to identify host factors involved in MVA infection. Deep-sequencing of gene trap insertion sites in the virus-resistant population identified genes involved in HepS biosynthesis, Golgi organization, vesicular protein trafficking and ubiquitination. Besides confirming TM9SF2 as a player in heparan sulfate biosynthesis, we identified TMED10 as a crucial factor for vaccinia virus-induced macropinocytosis.

## Materials and Methods

### Cells and viruses

The human melanoma cell line MelJuSo (MJS) and T2 cells were cultured in RPMI 1640 supplemented with 10% FCS, 100 U/ml penicillin, 100 μg/ml streptomycin and 2mM L-glutamine (complete medium). HEK-293T and Hela cells were cultured in DMEM supplemented with 10% FCS, 100U/ml penicillin, 100 μg/ml streptomycin and 2mM l-glutamine.

VACV strain Western Reserve (WR) encoding eGFP under control of the early/late P7.5 promoter (VACV-eGFP) was a generous gift from Dr. Jon Yewdell (NIH, Bethesda, USA). VACV-eGFP was propagated and titrated on Vero cells.

Recombinant modified vaccinia virus Ankara (MVA) expressing eGFP under the early/late promoter P7.5 (MVA-eGFP) or P11 late promoter (MVA-eGFP^late^) was propagated and titrated in chicken embryonic fibroblasts (CEFs). All viruses have been amplified, purified (sucrose cushion), and titrated according to standard methodology(19).

### Insertional mutagenesis

Mutagenesis of HAP1 cells using a retroviral gene trap vector was performed as described previously (20). For the screen, 1×10^8^ cells were infected with MVA-eGFP (MOI 50), and surviving cells were expanded and harvested for genomic DNA isolation. Retroviral insertion sites were amplified using a linear amplification-mediated PCR and deep-sequenced (Illumna HiSeq 2000). Sequences were aligned to the human genome (hg19) to identify retroviral insertion sites, which were assigned to non-overlapping protein-coding Refseq genes. Because the gene trap cassette was designed unidirectionally, genes functioning as MVA host factors would be predicted to be enriched for disruptive orientation insertions (21), as tested by a binomial test and corrected for multiple testing (Benjamini and Hochberg FDR). Genes already enriched in an unselected wildtype HAP1 dataset (NCBI Sequence Read Archive accession no. SRX1045466) were discarded.

### Lentiviral vectors

The selectable lentiviral CRISPR/Cas vector used in this study was previously described (22). This vector contains a human codon-optimized *S. pyogenes* Cas9 gene including a nuclear localization signal (NLS) at the N- and C-terminus. At the N-terminus, Cas9 is fused to PuroR via a T2A ribosome-skipping sequence under control of the human EF1A promoter. Additionally, it contains a human U6 promoter which drives expression of a guideRNA (gRNA) consisting of an 18-20bp target-specific CRISPR RNA (crRNA) fused to the *trans-*activating crRNA (tracrRNA) and a terminator sequence. This vector is called pSicoR-CRISPR-PuroR and has been described previously (22).

The crRNA target sequences for each of the targeted genes were designed using an online CRISPR design tool (crispr.mit.edu; Zhang lab, MIT). CRISPR gRNAs with the highest specificity and lowest off-target rate for the human genome were selected and cloned into the pSicoR-CRISPR-PuroR vector using Gibson assembly (NEB). The CRISPR gRNA-targeting sequences used in this study are listed in table 1.

**Table 1.**
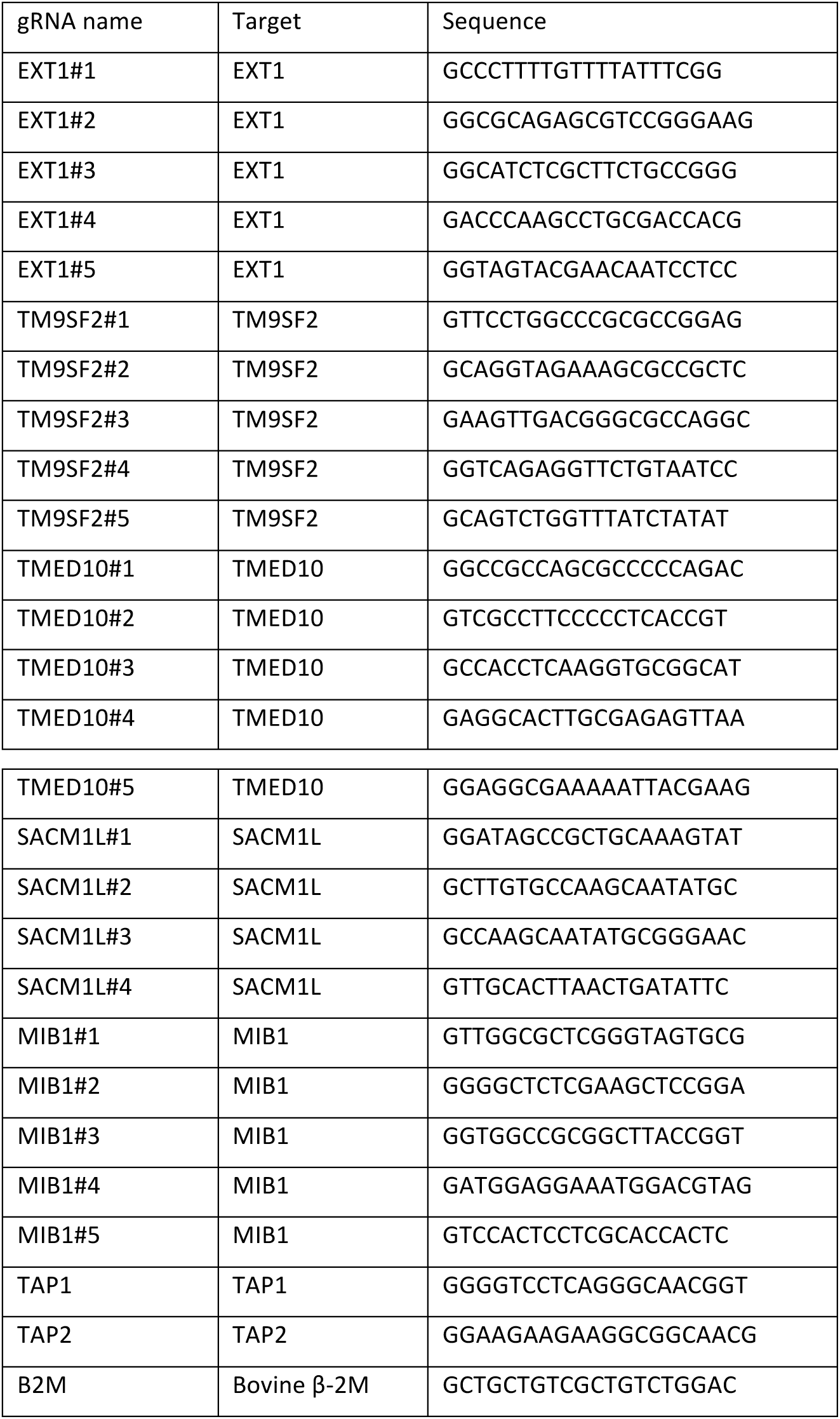
CRISPR gRNAs used in this study

For TMED10 rescue experiments, a gBLOCK (Integrated DNA technologies) containing the sequence of human TMED10 was introduced into a dual promoter lentiviral vector co-expressing ZeoR, and the gene encoding the fluorescent marker mAmetrine by means of Gibson assembly (NEB).

### Lentivirus production and transduction

Third generation lentiviruses were produced in HEK-293T cells in a 24 well plate format using standard lentivirus production protocols. MJS cells were transduced using spin infection at 1,000 x g for 90 minutes at 33°C in the presence of 3.2 μg/ml polybrene. After 3 days, transduced cells were selected using zeocin (400 μg/ml).

### Generation of knockout cell lines

MJS cells or HeLa cells were transfected with pSicoR_CRISPR_PuroR encoding the indicated gRNA (table 1), and subsequently selected using puromycin for 2 days (2 μg/ml). Clonal lines were generated by limited dilution after selection.

### Virus infections

For the validation in polyclonal MJS cell lines, 2 × 10^4^ cells/well were seeded in a 48 wells plate and infected the following day with MVA-eGFP using an MOI of 50. After seven days of infection, cells were harvested and the amount of cells was quantified by flow cytometry. Prior to flow cytometric analysis, 3,300 mCherry-positive T2 cells were mixed in to allow for normalization for cell counts between samples.

For infections in clonal Hela and MJS cells, 2,5 × 10^4^ cells were seeded in a 48 wells plate and infected the following day with the indicated virus strain at the MOI indicated. Cells were harvested five hours after infection with VAVC-eGFP and MVA-eGFP, and cells were harvested 20 h after infection with MVA eGFP^late^ to quantify the amount of infected cells (eGFP-positive) by flow cytometry.

### Heparan sulfate surface expression

Cells were harvested using an enzyme-free dissociation buffer (Sigma), and 5 × 10^4^ cells were incubated with the HepS-specific His-tagged monoclonal antibody (mAb) EV3C3 in PBS supplemented with 0.5% BSA and 0.02% sodium azide (PBA). Cells were washed with PBA and incubated with a FITC-labeled His-tag-specific mAb (AD1.1.10; Genxbio) in PBA. Cells were washed twice with PBA and fluorescence was quantified by flow cytometry (BD Biosciences) and analyzed by FlowJo (Treestar).

### SDS-PAGE and Western Blot analysis

Cells were lysed in 1% Triton buffer (1% Triton X-100, 20 mM 2- (N-morpholino) ethanesulfonic acid [MES], 100 mM NaCl, 30 mM Tris [pH7.5]) in the presence of 10 mM leupeptin and 1 mM 4- (2-aminoethyl) benzenesulfonyl fluoride. To remove nuclear fractions, lysates were centrifuged for at 12,000 x g at 4 °C for 20 min. To prepare samples for SDS-PAGE separation, postnuclear lysates were boiled in Laemmli sample buffer for 10min at 70 °C. After SDS-PAGE on Bolt 4-12% Bis Tris gels (Life Technologies), proteins were transferred to polyvinylidine fluoride membranes using the Trans-Blot Turbo Transfer system (Bio-Rad). Membranes were incubated with primary mouse antibodies specific for transferrin receptor (H68.4; Invitrogen), or TMED10 (clone A7; SantaCruz), washed, and incubated with the secondary HRP-conjugated goat anti-mouse IgG L-chain-specific antibody (Jackson ImmunoResearch #115-035-174). Bound antibodies were visualized by incubating membranes with ECL (Thermo Scientific Pierce) and exposure to Amersham Hyperfilm (GE Healthcare).

### Virus binding assay

Cells were harvested using enzyme-free dissociation buffer and 0.5 × 10^5^ cells were incubated with VACV-WR (MOI 20) in ice-cold complete medium. After 1h hour on ice, cells were washed in PBS supplemented with 10% FCS and fixed in PBS supplemented with 4% formaldehyde. Cells were incubated with an H3-specific polyclonal rabbit antibody, washed in PBA and incubated with a PE-conjugated goat anti-rabbit antibody. Cells were washed twice and fluorescence was quantified by flow cytometry.

### Virus blebbing assay

MJS cells (5 × 10^4^/well) were seeded on a Nunc Lab-Tek II Chamber Slide System (Thermo Scientific). The following day, VACV-WR was added (MOI 100), and cells were incubated on ice for 1 h to allow synchronized infection. Cells were subsequently incubated for 45 min at 37 °C, washed and fixed for 10 min at RT using PBS supplemented with 4 % formaldehyde. Cells were permeabilized in PBS supplemented with 0.1 % Triton for 10 min at RT, and subsequently incubated for 30 min with PBS supplemented with 0.5% BSA. Next, slides were incubated with CytoPainter phalloidin-iFluor 488 (Abcam) to stain actin filaments and TO-PRO3 (Thermo Fisher Scientific) to stain the nuclear DNA. Slides were washed and mounted on a microscope slide using Mowiol 4-88 (Carl Roth, Germany). Samples were imaged using a Leica TCS SP5 confocal microscope equipped with a HCX PL APO CS x 63/1.40-0.60 OIL objective (Leica Microsystems, the Netherlands). Fluorescent signals were detected with PMTs set at the appropriate bandwidth using the 488 nm argon laser for phalloidin and the 633 nm helium neon laser for TO-PRO3. Images were processed using the Leica SP5 software.

### Generation of Large Unilamellar Vesicles

Calcein-encapsulated LUVs were composed of 1,2-dioleoyl-*sn*-glycero-3-phosphocholine (DOPC) or a mixture of DOPC and 1,2-dioleoyl-*sn*-glycero-3-phospho-L-serine (DOPS) (Avanti Polar Lipids) in a 7:3 molar ratio. Stock solution of DOPC and DOPS in chloroform (10 mM) were mixed in a glass tube, and chloroform was subsequently evaporated with dry nitrogen gas. The yielded lipid film was subsequently kept in a vacuum desiccator for 20 min. Lipid films were hydrated for 30 min in buffer containing 10 mM Tris, 50mM NaCl (pH 7.4) resulting in total lipid concentration of 10 mM. For calcein-encapsulated LUVs, 50 mM of calcein was added during hydration. The lipid suspensions were freeze-thawed for 10 cycles, at temperatures of −80 and +40°C, and eventually extruded 10 times through 0.2 μM-pore size filters (Anotop 10, Whatman, UK). Free calcein was separated from calcein-filled LUVs using size exclusion column chromatography (Sephadex G-50 fine) and eluted with 10 mM Tris-HCl, 150mM NaCl buffer (pH 7.4). The phospholipid content of lipid stock solutions and vesicle preparations were determined as inorganic phosphate according to Rouser (23). Finally, average LUV size of 150-200 nm was confirmed using dynamic light scattering.

### LUV *macropinocytosis*

Cells were seeded in a 48 wells plate using 5 × 10^4^ cells per well. The following day, cells were incubated for 30 min with EIPA where indicated and subsequently incubated with LUVs for 40 min in complete medium or medium lacking serum where indicated. Cells were subsequently harvested, and washed twice in ice-cold PBS. Cells were fixed in PBS supplemented with 1% formaldehyde, and fluorescent calcein signal was quantified by flow cytometry. For microscopic analysis of LUV uptake, cells were seeded onto a Nunc Lab-Tek II Chambered coverglass (Thermo Fisher Scientific). The next day, cells were incubated with LUVs and directly imaged in a climate chamber set at 37 °C using the bright field camera and mercury –vapor lamp of the Leica SP5 confocal microscope.

## Results

### A haploid genetic screen identifies host factors for MVA infection

To identify host factors involved in MVA infection, we performed a genome-wide haploid genetic screen using HAP1 cells, which are readily infected and ultimately killed by MVA. A total of 1 × 10^8^ HAP1 cells were mutagenized using a gene trap retrovirus and subsequently infected with MVA. Resistant HAP1 cells were propagated for ten days and their retroviral insertion sites were subsequently identified by deep sequencing to recognize the disrupted genes. Genes that were enriched for disruptive gene trap mutations in the virus selected population but not in uninfected control cells were identified (fig. 1 and table S1). Stongest outliers were host factors that directly affect HepS chain formation, including XYLT2, B4GALT7, B3GALT6, B3GAT3, EXTL3, EXT1, EXT2, HS2ST1, and NDST1 (reviewed by (24)). Other host factors affecting HepS transport and biosynthesis of HepS precursor molecules were also enriched in the screen, including UGDH, UXS1 SLC35B2, and PTAR1 (20, 21, 25-28). A second cluster of host factors identified in the screen included all the subunits of the Conserved Oligomeric Golgi (COG) complex. This complex is involved in the distribution of glycosylation enzymes in the Golgi (29), and thereby indirectly affects HepS expression (20). Several other significant hits did not cluster with other genes for functional relationship, and were involved in multiple cellular processes, including protein and vesicle trafficking, ubiquitination, and Golgi organization.

**Figure 1.**
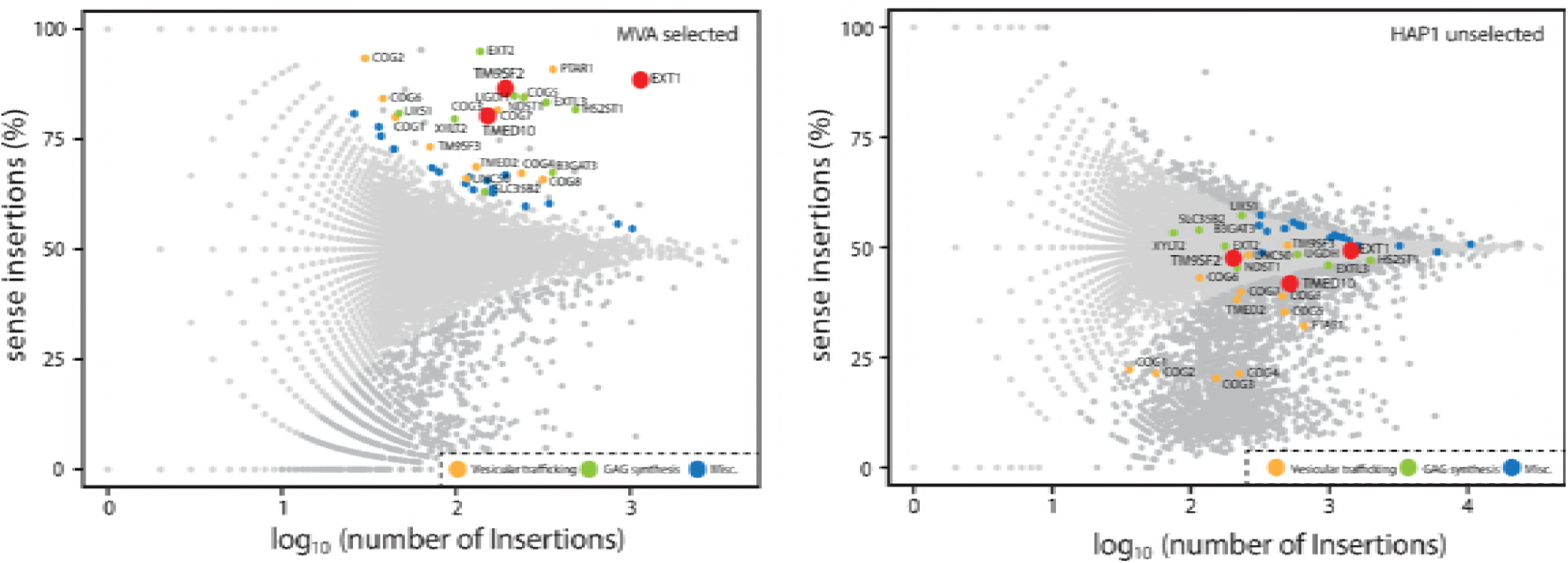
Genome-wide haploid genetic screen identifies host factors required for MVA infection. Genes enriched for retroviral insertion sites in MVA-exposed cells (left panel) compared to an unexposed control population (right panel). The percentage of sense orientation gene-trap insertions is plotted on the y axis. The total number of insertions in a particular gene is plotted on the x axis. Genes indicated by larger red dots were further characterized in this study.

### Validation of hits using the CRISPR/Cas9 system

We used the CRISPR/Cas9 system to validate the a number of hits, including EXT1, TM9SF2, TMED10, and SACM1L. These genes were selected based on the significance of the enrichment and biological interest. In addition, mindbomb protein 1 (MIB1) was selected as a hit that was already significantly enriched in wild type HAP1 cells (see table S1). Up to five gRNAs per gene were selected and co-expressed with Cas9 in the human melanoma cell line MelJuSo (MJS), which was readily infected by the poxviruses used in this study. In addition, control gRNAs were expressed that target TAP1, TAP2, or B2M, which have no known role in primary poxvirus infection. Cells were infected with MVA at an MOI of 50, and surviving cells were counted seven days post infection. The majority of the wild type MJS cells and cells expressing control gRNAs were susceptible to virus-induced cell death. In contrast, four out of the five gRNAs targeting EXT1 and TM9SF2 conferred robust (≥50%) protection from MVA-induced cell death (fig. 2). Similarly, five out of the five gRNAs targeting TMED10 protected the majority of the cells, although not as pronounced as for gRNAs targeting EXT1 or TM9SF2. In contrast, only one gRNA targeting SACM1L and no gRNAs targeting MIB1 conferred robust protection against MVA infection. Thus, gRNAs targeting EXT1, TM9SF2, and TMED10 protected cells from virus-induced cell death.

**Figure 2.**
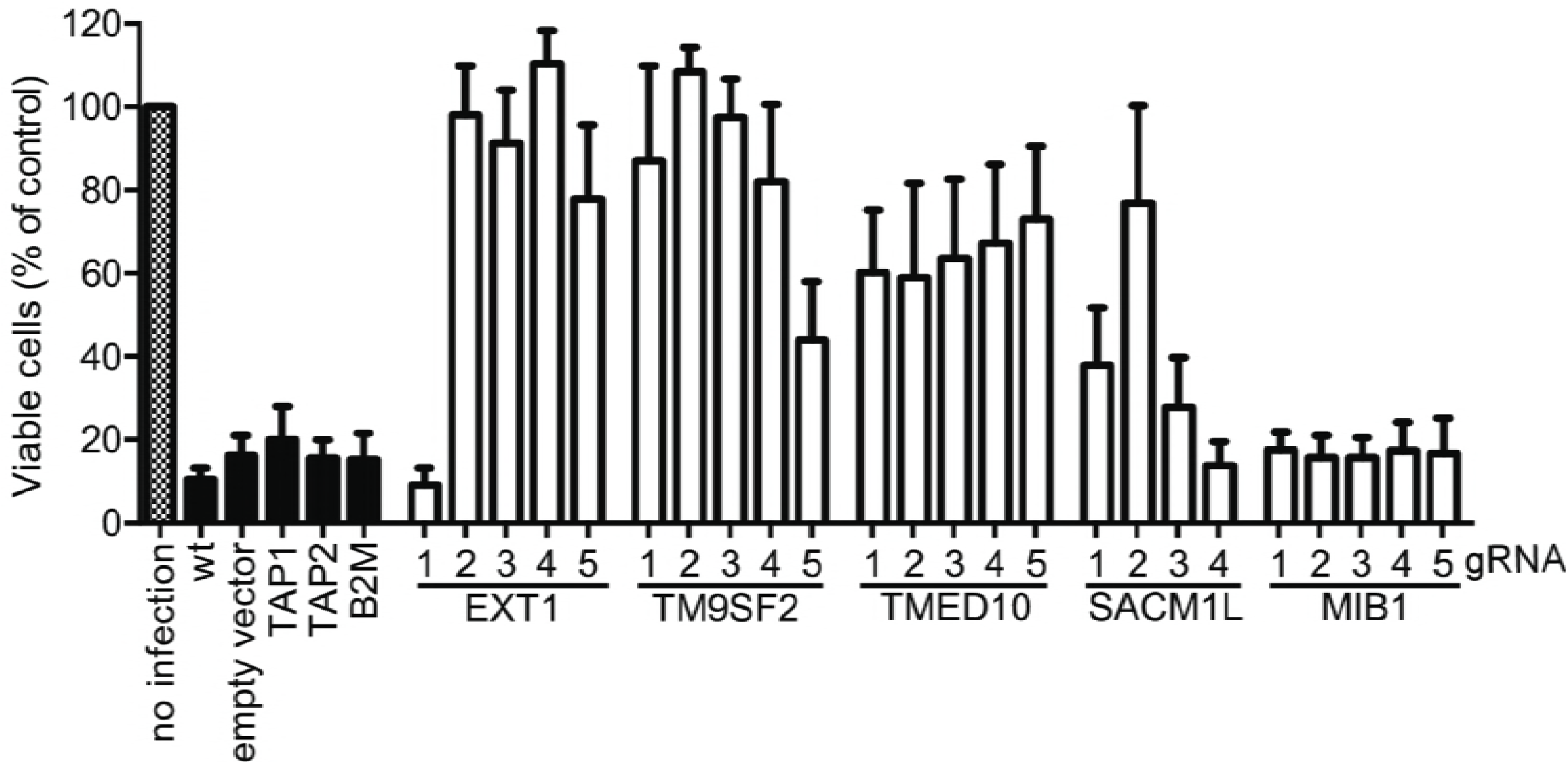
EXT1, TM9SF2, TMED10, and SACM1L are essential genes for MVA infection. Validation of hits in MJS cells. Wild type MJS cells (wt) and MJS cells transfected with Cas9 and indicated gRNAs (see table 1) were infected with MVA-eGFP (MOI 50). After 7 days of infection, cells were harvested and quantified by flow cytometry. Data are represented as S.E.M. of three independent infection experiments.

### EXT1 and TM9SF2 affect MVA infection through HepS expression

Survival from MVA exposure may be due to resistance to virus-induced cell death, or resistance to primary virus infection. To assess their role in the primary infection event of MVA, anti-EXT1 and anti-TM9SF2 gRNA-expressing cells were cloned and subsequently infected with MVA expressing eGFP from an early/late promoter and monitored for eGFP expression 5h post infection (fig. 3A). Several, but not all, clones expressing EXT1 or TM9SF2 gRNAs were highly resistant to MVA infection. Primary infection was reduced more than 70% in four out of five EXT1 gRNA clones and more than 60% in four out of eleven TM9SF2 gRNA clones (fig. 3A).

**Figure 3.**
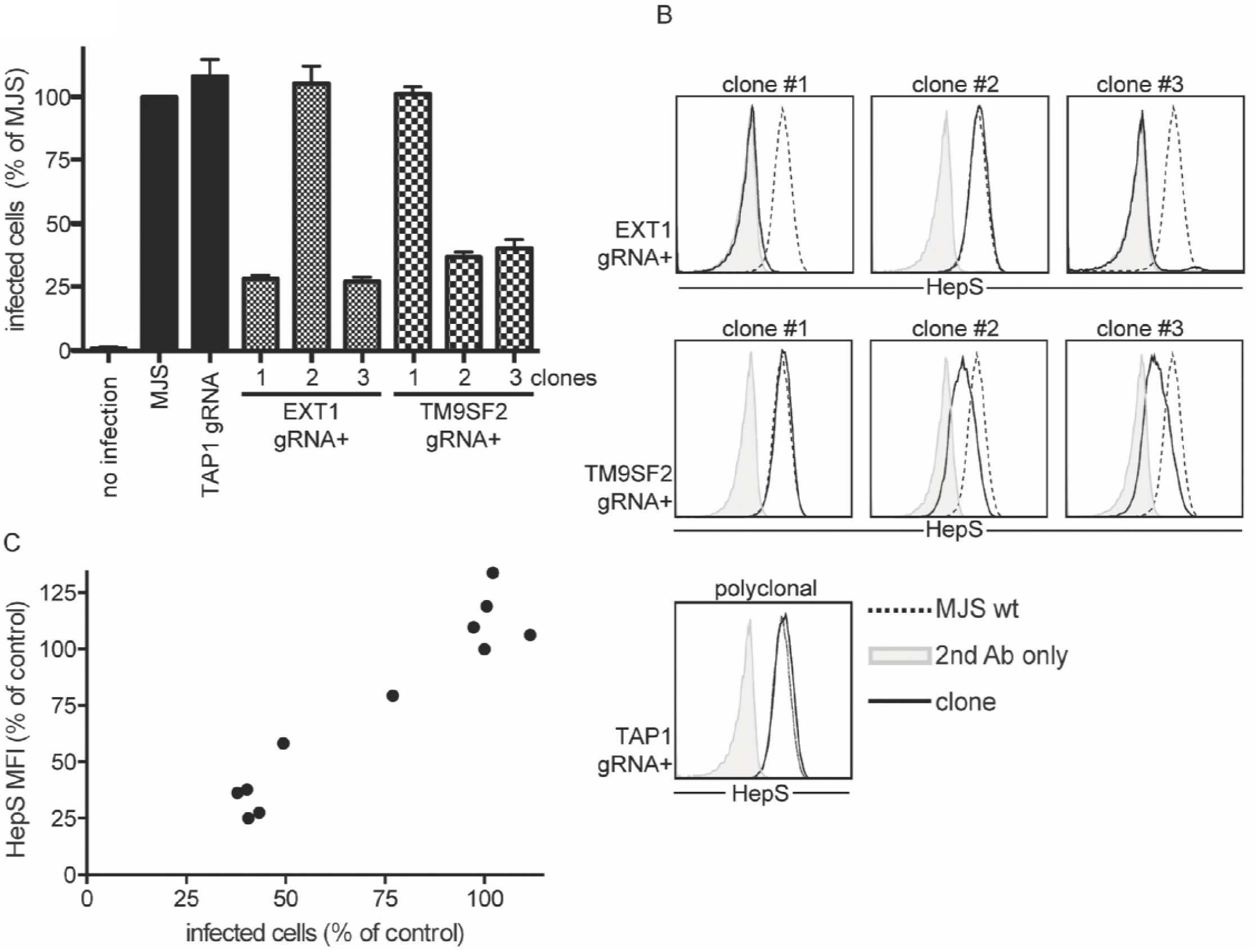
Efficiency of MVA infection depends on HepS surface levels. (A) Wild type MJS cells (wt) and MJS cells transfected with Cas9 and gRNA TAP1, EXT1#2, EXT1#4, TM9SF2#1 or TM9SF2#2 were cloned and infected with MVA-eGFP (MOI 10). After five hours of infection, cells were harvested and the amount of infected cells (GFP-positive) was quantified by flow cytometry. S.E.M. of three independent infections is indicated. Three representative clones of five are shown for EXT1, four of these clones were protected from MVA infection. Three representative clones of eleven are shown for TM9SF2. (B) Clonal lines indicated in (A) were analyzed for HepS expression levels by flow cytometry. (C) Eleven clonal lines obtained from MJS cells transfected with gRNA TM9SF2#1 or TM9SF2#2 were infected with MVA-eGFP (MOI 10). In addition, the cells were stained for HepS surface expression and the mean fluorescent intensity (MFI) was quantified by flow cytometry.

EXT1 is a crucial player in HepS synthesis by catalyzing the final steps in HepS chain formation. TM9SF2 appeared as a hit in a haploid screen for HepS surface expression (20), and was more recently identified as a crucial factor for N-deacetylase/N-sulfotransferase 1 (NDST1) localization and functioning(30). We tested the effect of the EXT1 and TM9SF2 gRNAs on HepS surface expression in the same clones presented in fig. 3a. In the EXT1 gRNA-expressing clones that were resistant to primary MVA infection, HepS surface expression was reduced to background levels. In contrast, EXT1 clone 2, which was not resistant to MVA infection, displayed unaltered HepS surface levels (fig. 3B). Similar results were obtained for the TM9SF2 clones, although the downregulation of HepS surface levels was less pronounced as compared to the EXT1 clones. A total of 11 TM9SF2 clones were tested for their resistance to MVA infection and HepS surface expression. These clones displayed varying levels of HepS surface expression, which correlated with MVA infection (fig. 3C). These results confirm a key role for EXT1 and TM9SF2 in HepS surface expression.

### TMED10 is necessary for an early stage of vaccinia virus infection

Next, we tested the role of TMED10 in primary MVA infection of HeLa and MJS cells. TMED10 gRNA-expressing HeLa and MJS cells were cloned and tested for TMED10 expression by immunoblotting (fig. 4A). Although we could readily knock-out TMED10 from MJS cells, we were not able to establish TMED10-null HeLa cells, suggesting that TMED10 is essential in HeLa but not MJS cells (fig. 4A). In line with this, the TMED10 knockdown clones we could establish in HeLa cells displayed decreased growth rate. In contrast, removing TMED10 from MJS cells did not result in altered morphology or growth rate.

**Figure 4.**
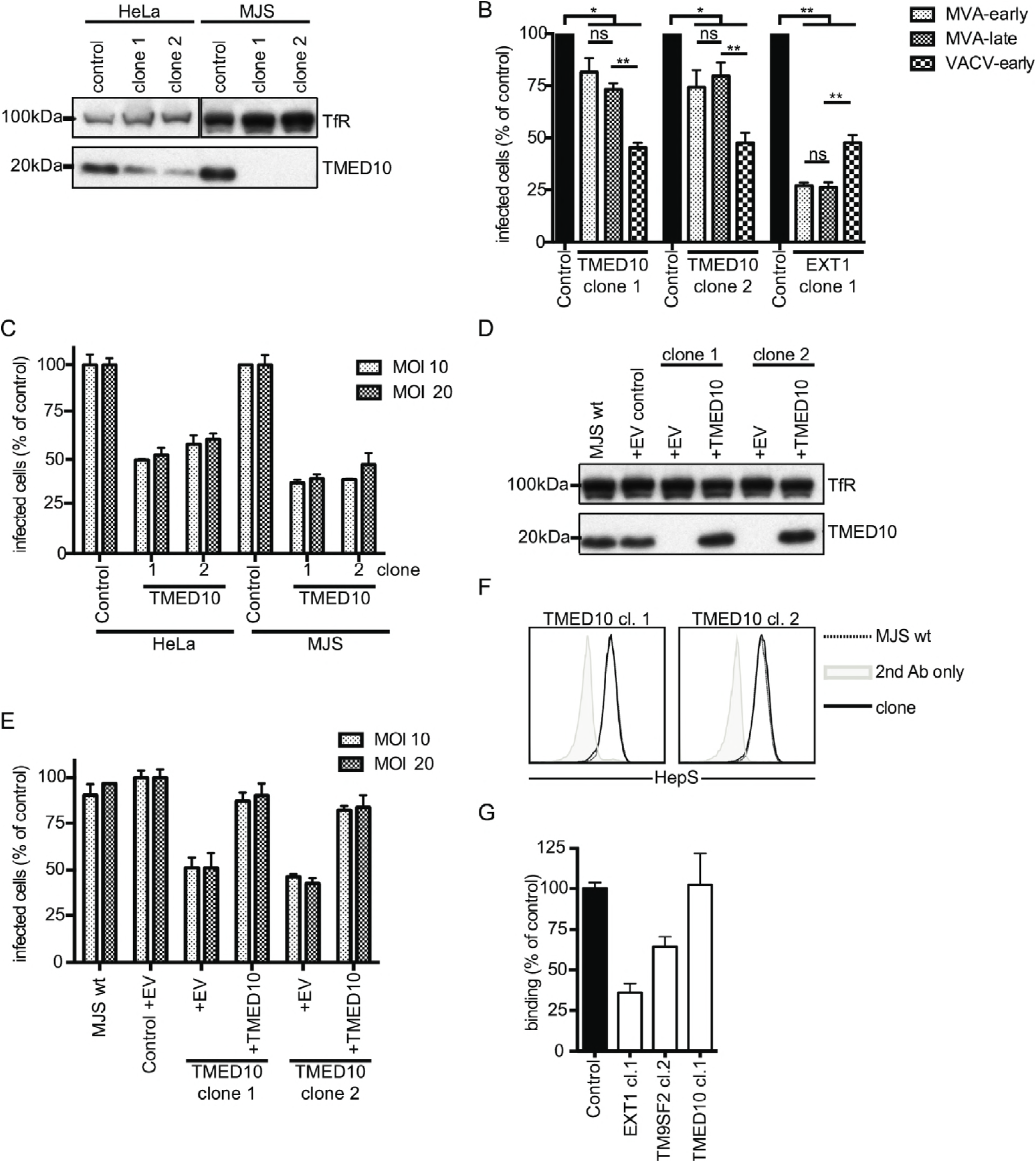
TMED10 restricts infection of MVA- and VACV-eGFP. (A) HeLa cells expressing Cas9 only or co-expressing TMED10#1 and TMED10#5, and MJS cells expressing Cas9 only or co-expressing TMED10#1 or TMED10#5 were cloned, and transferrin receptor and TMED10 levels were determined by immunoblotting. The immunoblot is representative of three independent experiments. (B) MJS cell clones indicated in (A) or EXT1 clone 1 (see fig. 3A) were infected with MVA-eGFP, MVA-eGFP^late^, or VACV-eGFP. After five hours, cells infected with MVA-eGFP, and VACV-eGFP were harvested to determine early promoter-driven eGFP expression of MVA-eGFP (MVA-early) or VACV-eGFP (VACV-early). After 20 hours of infection with MVA-eGFP^late^, cells were harvested to determine eGFP expression driven from a late promoter (MVA-late). S.E.M. of four independent experiments is shown. ns: not significant; *p<0.05; **p<0.0005. Significance was calculated using a two-tailed unpaired t-test. (C) MJS cells and HeLa cells indicated in (A) were infected with VACV-eGFP at either MOI of 10 or 20, and the percentage of infected (eGFP-positive) cells was quantified by flow cytometry. S.E.M. of three independent experiments is shown. (D) Wild type MJS cells or MJS clones indicated in (A) were transduced with an empty vector (+EV) lentivirus or a lentivirus encoding TMED10 (+TMED10). Transferrin receptor and TMED10 expression levels were assessed by immunoblotting. The immunoblot shown is representative of two independent experiments. (E) Wild type MJS cells or clones indicated in (D) were infected with VACV-eGFP. The percentage of infected (eGFP-positive) cells was quantified by flow cytometry five hours after infection. S.E.M. of four independent experiments is shown. (F) Flow cytometric analysis of wild type MJS (dashed line) and MJS TMED10 clones 1 and 2 (black lines) stained for HepS surface-expression, or secondary antibody only (gray filled line). (G) VACV-WR was allowed to bind control cells or indicated clones on ice. After one hour, cells were washed, fixed, and stained with an antibody specific for the viral H3 protein. Bound antibody was quantified by flow cytometry. The mean fluorescence intensity of the control cells was set at 100%. S.E.M. of three experiments is shown.

The role of TMED10 in primary infection was tested by infecting MJS TMED10 KO clones with two recombinant MVA strains that express eGFP during infection controlled by either an early/late or late promoter. To measure early promoter activity, eGFP expression was evaluated 5h after infection (fig. 4B). In both TMED10 KO clones, infection was reduced modestly but significantly compared to control cells. To test whether TMED10 knockout had an enhanced effect during late stages of infection, cells were infected with MVA expressing eGFP from a late promoter, and eGFP expression was monitored 24h after infection. Late gene expression was affected to similar levels as early gene expression, suggesting that TMED10 affects virus infection prior to early gene expression (fig. 4B). A similar reduction in early and late gene expression was also observed in MJS EXT1 clone 1, thereby confirming that EXT1 affects MVA infection before the onset of early gene expression (fig. 4B).

To test the role of TMED10 in infection with other vaccinia virus strains, MJS cells were infected with vaccinia virus strain Western Reserve (VACV-WR) expressing eGFP from an early/late promoter (VACV-eGFP). Five hours after inoculation, VACV-eGFP infection was significantly more abrogated as compared to the infection with MVA, and showed a reduction of over 50% when compared to control cells. VACV-WR infection was also highly reduced in MJS EXT1 clone 1, although not as pronounced as for MVA. This is in line with previous reports showing that VACV-WR is less dependent on HepS for attachment than the MVA strain (31, 32) (fig. 4B). Next, we tested the role of TMED10 in VACV-eGFP infection of HeLa cells. Despite the incomplete removal of TMED10 protein from these cells (fig. 4A), VACV-eGFP infection was reduced up to 50% in both HeLa TMED10 knockdown clones at differtent MOIs. In the two MJS TMED10 KO clones, infection was again highly reduced at both MOIs tested (fig. 4C).

To exclude that the observed phenotype was caused by off-targeting effects of the two TMED10 gRNAs, the TMED10 cDNA was reintroduced in the MJS TMED10 KO clones by means of lentiviral transduction (fig. 4D). Indeed, TMED10 reconstitution facilitated VACV-WR infection to similar levels as control cells (fig. 4E). Thus, we identified TMED10 as a new host factor that affects vaccinia virus infection before the onset of early gene expression. Similar to EXT1 and TM9SF2, TMED10 may affect HepS surface expression and thereby impact attachment of the virus to the host cell membrane. However, both TMED10 KO MJS clones displayed similar HepS surface levels as compared to wt MJS cells (fig. 4F). Furthermore, TMED10 KO had no effect on VACV-WR binding to the host cell membrane, in contrast to EXT1 and TM9SF2 KO cells (fig. 4G). To conclude, TMED10 affects poxvirus infection post attachment but prior early gene expression, suggesting a role during virus entry.

### TMED10 is crucial for membrane blebbing and macropinocytosis

The more pronounced effect of TMED10 on VACV-eGFP infection compared to MVA infection may result from the differences in entry pathways that both viruses use. Whereas MVA mainly enters cells by virus-cell fusion at the plasma membrane, VACV predominantly enters the cell through macropinocytotic uptake of virus particles (33). Macropinocytosis of vaccinia is a well-described process that is dependent on the presence of phosphatidylserine (PS) in the viral envelope (34, 35). Interactions between PS and PS receptors are facilitated by serum factors, and induce changes in membrane morphology (36-39). These actin-based changes lead to the formation of giant blebs on the surface of affected cells. Upon retraction of the bleb, virus particles are enclosed in a newly formed macropinosome, which subsequently moves further into the cytoplasm (35).

The role of TMED10 in the formation of virus-induced blebs was investigated by incubating control cells and TMED10 KO cells with VACV-WR for 45 min. Subsequently, actin filaments were visualized using confocal microscopy. In control cells, blebbing was readily observed on the surface of virus-exposed cells, whereas blebs were absent in uninfected cells (fig. 5). In contrast, virus-induced blebbing was not observed in TMED10 KO cells, although reintroduction of TMED10 restored blebbing in these cells (fig. 5).

**Figure 5.**
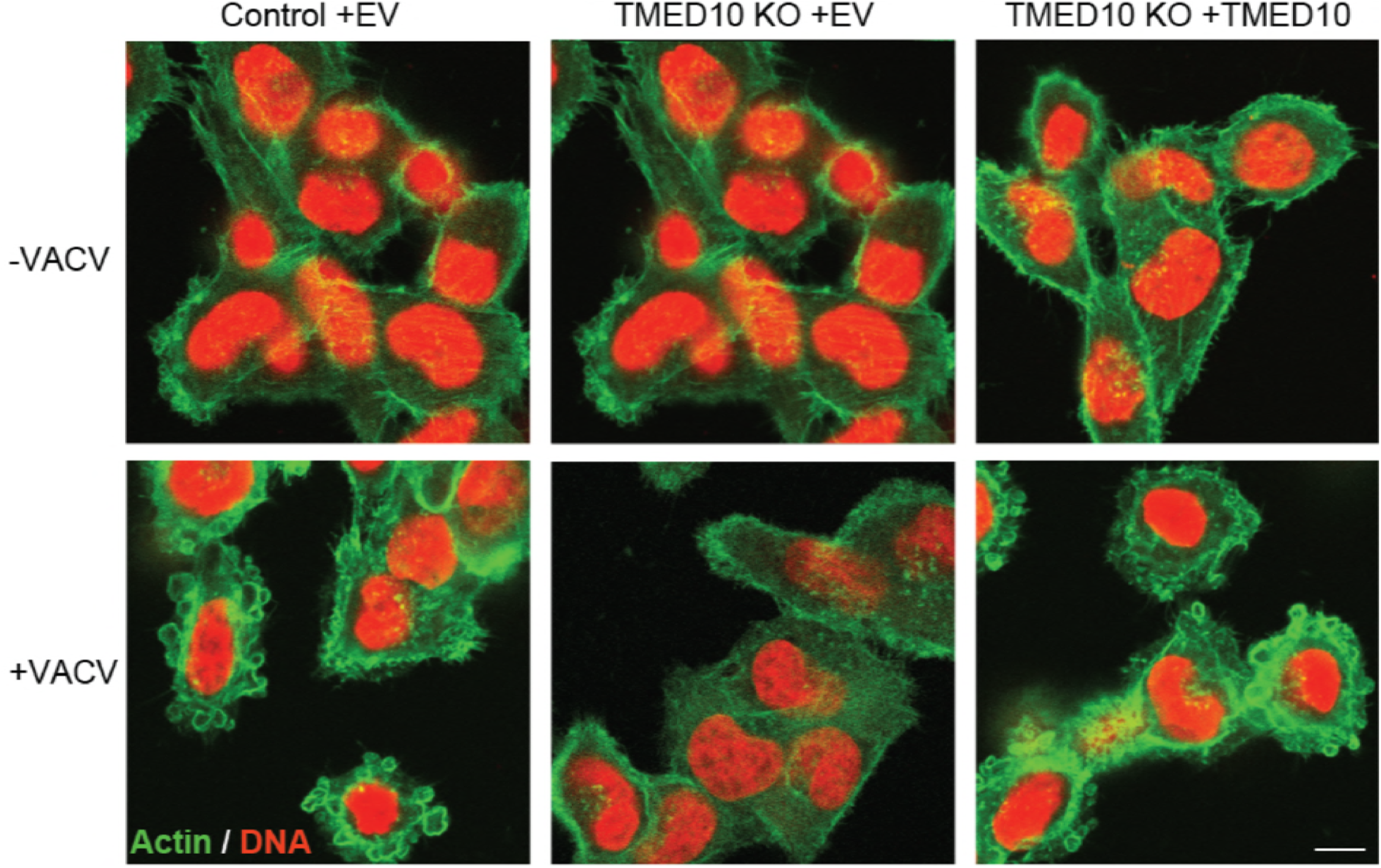
Virus-induced formation of blebs is dependent on TMED10 expression. Control MJS cells expressing an empty vector (EV) or TMED10 clone 2 expressing an EV or TMED10 (see fig. 4D) were incubated with VACV-WR (MOI 100) (+VACV) or left untreated (-VACV) for one hour on ice. Subsequently, the cells were incubated at 37C for 45 min to allow bleb formation. Blebs were visualized by staining the cells for actin (green) using phalloidin iFluor 488; nuclei were counterstained (red) using TO-PRO3. A representative image field of 20 different image fields is presented. Bar size: 7.5 μm.

The role of TMED10 in macropinocytosis was further demonstrated using large unilamellar membrane vesicles (LUVs). These LUVs are similar in size (150-200 nm in diameter) as vaccinia viruses and can be taken up by cells via macropinocytosis (fig. S1). By introducing the self-quencing fluorescent marker calcein in LUVs, their uptake in cells can be assessed as intracellular LUV release results in the formation of a bright fluorescent signal in the cell (fig. S1). As for VACV, the concentration-dependent uptake of LUVs is dependent on serum factors and PS in the membrane (fig. S1B). In addition, LUV uptake can be blocked by the macropinocytosis inhibitor EIPA (fig. S1B). LUV uptake was quantified by flow cytometry in control cells and cells lacking TMED10 (fig. 6). In TMED10 knockdown HeLa cells and TMED10 KO MJS cells LUV uptake was highly reduced, as quantified by flow cytometry (fig. 6A). LUV uptake was restored upon reintroduction of TMED10 (fig. 6B). In conclusion, these results indicate that TMED10 affects virus-associated membrane blebbing and the subsequent uptake of virus-like particles through PS-dependent macropinocytosois.

**Figure 6.**
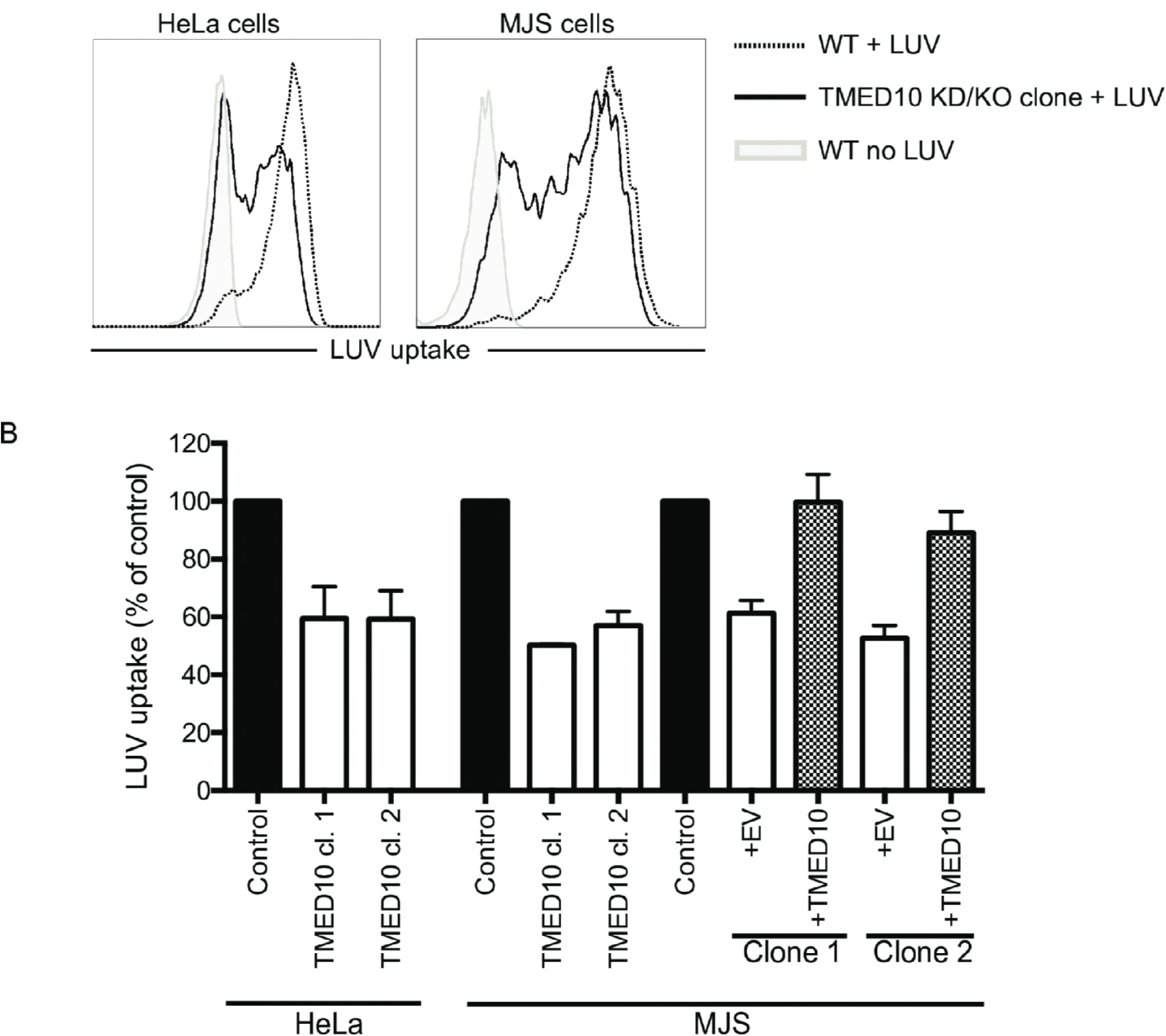
Macropinocytotic uptake in Hela and MJS cells is dependent on TMED10. A) Flow cytometric analysis of control Hela and MJS cells (dashed lines), Hela TMED10 clone 1 and MJS TMED10 clone 1 (black lines) incubated for 40 min at 37°C degrees with 3 μM fluorescent LUVs composed of PC and PS. Cells were harvested and fluorescence was determined by flow cytometry. (B) MFI of LUV uptake was measured as in (A) in the indicated Hela and MJS clones. LUV uptake was normalized to wt cells incubated with LUV. S.E.M. of four independent experiments is shown.

## Discussion

This report represents the first genome-wide haploid genetic screen that aimed to identify host factors important for vaccinia virus infection. The relative absence of host factors identified in this screen at later stages of MVA infection may in part be due to the use of replication-incompetent virus. MVA infection is abrogated in most cell lines due to defects in virus assembly (40). Therefore, host factors involved in later stages of infection are not selected for in a given screen. Screening approaches using replication-competent vaccinia viruses may elucidate more of these factors, although our initial results showed that such viruses are too aggressive and did not allow recovery of virus-resistant cells. Another factor that may contribute to the bias towards host factors affecting early stages of virus infection is the selection method of this screen. This is based on resistance to virus-induced cell death, and thereby does not account for selection of genes mediating virus production and spread. In line with this, a haploid screen using replication-competent RVFV or pseudotyped VSV also recovered host factors mostly involved in virus attachment and entry (20, 27, 41, 42).

This study clearly illustrates the dominant role of HepS in vaccinia virus infection. This is in line with previous studies showing that attachment of MVA and other vaccinia strains to the host cell is primarily mediated by interactions with HepS on the cell surface (43-46). Although MVA and VACV-WR infection is abrogated in the absence of HepS surface expression, a proportion of the cells can still become infected (see fig. 3A and 4B). It is suggestive that attachment of viruses in the absence of HepS is mediated by other glycosaminoglycans, including chondroitin sulfate and dermatan sulfate (47). Other factors including the extracellular matrix protein laminin may also participate in virus attachment(48). In addition, unidentified cell surface receptors have been implicated in GAG-independent binding to the cell (15). Although our study did not reveal a specific MVA attachment mediator, the preeminent role of HepS may mask the role for such alternative entry mechanisms. In that respect, additional host factors could be identified by performing a screen on HepS-deficient HAP1 cells.

The HepS-associated host factors identified in this screen largely overlap with factors identified in haploid genetic screens performed for other viruses that (partly) depend on HepS for host cell attachment, including Lassa virus, RVFV, and adeno-associated virus (27, 42, 49). In addition, a haploid genetic screen that directly assessed the role of host factors involved in HepS biosynthesis revealed a similar enrichment of genes and also identified EXT1 as the most significant hit (42). Other genes that were enriched in these screens included TM9SF2 and prenyltransferase alpha subunit repeat containing 1 (PTAR1). The involvement of PTAR1 in HepS surface expression has been confirmed recently (20, 21, 27). TM9SF2 is a member of the highly conserved nonaspanin proteins that have been connected to diverse cellular processes, most notably receptor trafficking, and cellular adhesion (50, 51). In addition, TM9SF2 affects the localization and stability of NDST1, which is critical for N-sulfation of HepS (30). TM9SF2 single nucleotide polymorphisms have been associated with several human diseases, including progression of AIDS(52). This association may, in part, be explained by a TM9SF2-mediated decrease of HepS, thereby affecting binding of HIV to the cell surface (53).

We identified TMED10 (also known as TMP21/p23/p24δ) as an important host factor for VACV infection. TMED10 is a ubiquitously expressed type I transmembrane protein that associates with coat complex protein I (COPI) vesicles through KKLIE motif in its cytosolic tail (54, 55). As such, it is suggested to function as a cargo receptor in retrograde vesicular trafficking from the Golgi to the ER (54, 56, 57). In addition, TMED10 localizes to the plasma membrane independently of COPI (58). Plasma membrane-localized TMED10 controls the activity of the presenilin/γ -secretase complex in neuronal tissue, thereby affecting the formation of amyloid beta peptides implicated in Alzheimer’s disease (59). Other binding partners have also been described for TMED10, including ER-localized MHC class I (60), and C1 domain-containing proteins such as the chimaerins (61, 62). These latter proteins are critical regulators of the actin cytoskeleton modulator Rac1 (61-64).

The regulation of chimaerins by TMED10 may explain the effects on virus-induced macropincotytosis observed in this study. Vaccinia-induced blebbing and subsequent macropinocytosis critically depend on actin rearrangements regulated by the Rho GTPase Rac1 (34, 35, 37, 65, 66). Rac1 is deactivated by β2-chimaerin localized at the plasma membrane (61). This localization is regulated by TMED10 that redistributes β2-chimaerin to the perinucleus upon binding (61, 62) Conversely, depletion of TMED10 or disruption of the β2-chimaerin/TMED10 complex relocates β2-chimaerin to the plasma membrane, and thereby enhances Rac1 deactivation (61). Similarly, Rac1 deactivation in TMED10 KO cells could explain the inhibitory effect on blebbing and macropinocytosis observed in this study. We identified TMED10 as an important factor in PS-induced macropinocytosis. This pathway is not only used by vaccinia viruses to enter cells; many other viruses expose PS to allow entry by macropinocytosis (reviewed in (34)). In contrast to plasma membrane fusion, macropinocytosis allows viruses to bypass the dense cortical actin layer to enter the cytosol. Macropinocytosis also aids in immune escape of viruses, as this uptake pathway minimizes exposure of viral antigens on the cell surface (16). In addition, viruses may benefit from the dampening effect of PS-induced macropinocytosis on the immune system, which is mediated by an array of anti-inflammatory cytokines (34, 67, 68). It would be interesting to know if TMED10 is also vital for entry of other viruses, thereby facilitating their immune escape.

## Author Contributions

Conceptualization, R.D.L, R.J.L., and E.J.W; Methodology, V.A.B., and T.R.B.; Investigation, R.D.L., F.V.D., and V.A.B.; Writing – Original Draft, R.D.L.; Writing –Review & Editing, R.D.L., V.A.B., T.R.B., R.J.L., and E. J.W.; Resources, S.M.S., I.D., and T.H.V.K.; Supervision, T.R.B., R.J.L., and E.J.W.

## Acknowledgements

We thank Ronny Tao for technical assistance. RJL was supported by Marie Curie Career Integration Grant PCIG-GA-2011-294196. SMS was supported by the seventh framework program of the European Union (Initial Training Network “ManiFold,” Grant 317371), and ID was supported by the DFG funding GRK 1949. The authors declare that they have no conflict of interest.

## Data availability

Data will be made available on public servers prior to publication

---

**Table S1.** List of genes significantly enriched in MVA-selected HAP1 cells.

**Figure S1.**
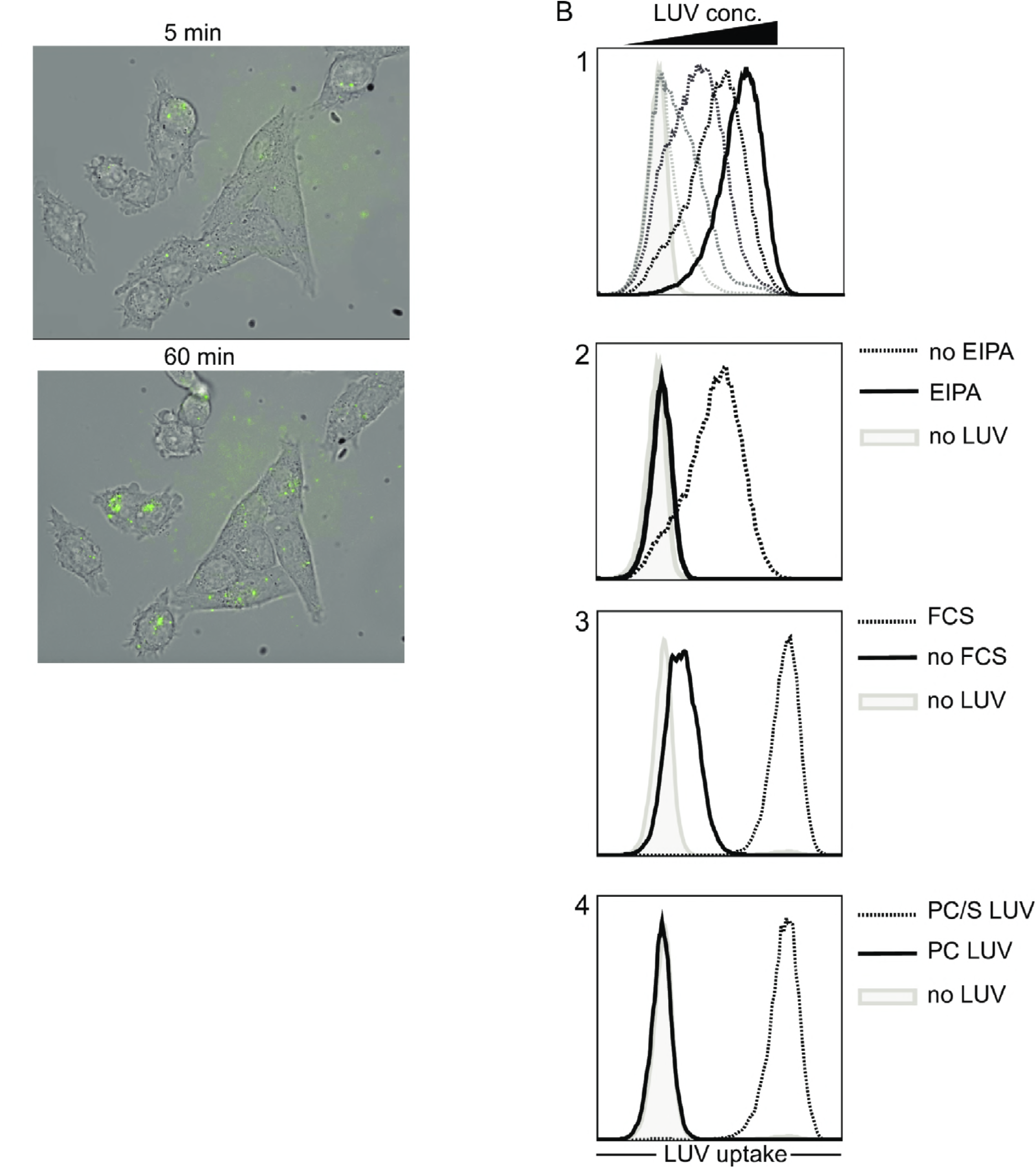
Uptake of LUVs is mediated by macropinocytosis and depends on PS and serum components. (A) Microscopic tracing of cells exposed to 3 μM fluorescent LUVs composed of PC and PS. (B) Flow cytometric analysis of MJS cells incubated with fluorescent LUVs for 45 min. Panel 1: uptake using different concentrations of LUVs (6 to 0.2 μM). Panel 2: LUV uptake (3 μΜ) in the absence (dashed line) or presence or the specific macropinocytosis inhibitor EIPA (black line). Panel 3: LUV uptake (3 μΜ) in the presence (dashed line) or absence (black line) of serum. Panel 4: uptake of LUVs (3 μΜ) composed of PC only (black line) or PC and PS (dashed line). Gray filled histograms: no LUVs added.

